# Evidence of climate change impacts on the iconic *Welwitschia mirabilis* in the Namib Desert

**DOI:** 10.1101/2020.02.19.955823

**Authors:** Pierluigi Bombi, Daniele Salvi, Titus Shuuya, Leonardo Vignoli, Theo Wassenaar

## Abstract

Climate change represents an important threat to global biodiversity and African ecosystems are particularly vulnerable. Recent studies predicted substantial variations of climatic suitability for *Welwitschia mirabilis* under future conditions. Latitudinal/altitudinal range shifts are well-known responses to climate change but not coherent patterns were documented. This study aims to verify whether welwitschia populations are responding to climate change and if the assumption of a latitudinal/altitudinal shift is applicable. We collected field data on welwitschia distribution, health condition, reproductive status, and plant size in northern Namibia. We used ecological niche models to predict the expected geographic shift of climatic suitability under future scenarios. For each variable, we compared the observed pattern with the expected responses. Finally, we tested the presence of simple geographical gradients in the observed patterns. The realized thermal niche of welwitschia will be almost completely unavailable in the next 30 years in northern Namibia. Expected reductions of climatic suitability in the stand sites are strongly associated with indicators of negative population conditions. The same population conditions does not fit any simple latitudinal or altitudinal gradient. The observed pattern of population conditions mirrors the expected pattern of climate change effect but no simple geographical gradient was relieved. Overall, we observed negative population conditions in areas with stronger reductions of suitability. This makes welwitschia a suitable sentinel for climate change effect in the Namib Desert ecosystems. Our approach to detect population responses to climate change could be extensively adopted for selecting sentinel species in other regions and ecosystems.

## 1. Introduction

Climate change is one of the strongest threats for ecosystems worldwide. Variations in the density of species, range shifts, and extinction events have been documented at local and global level (Cristofari et al., 2018; Parmesan and Yohe, 2003; Walther et al., 2002). Furthermore, changes in species diversity, ecosystem functioning, and service provision are expected for the future as a consequence of additional pressures on natural populations (Ge et al., 2015; e.g. Hole et al., 2009; Moritz and Agudo, 2013). In Africa, deep impacts by climate change were forecasted for animals (e.g. Garcia et al., 2012; Huntley and Barnard, 2012; Kirchhof et al., 2017), plants (Blach-Overgaard et al., 2015; e.g. Midgley et al., 2003; Midgley and Bond, 2015), and biodiversity conservation in general (Hole et al., 2009; Revermann et al., 2018). Arid regions of southern Africa seem to be particularly exposed to the effects of climate change (Midgley and Thuiller, 2011). For the quiver tree (*Aloidendron dichotomum* Klopper & Gideon 2013), climate-linked increases of mortality were observed, although this evidence is still controversial (Foden, 2002; Foden et al., 2007; Guo et al., 2011; Jack et al., 2016).

Recently, Bombi (2018) highlighted potential effects of climate change on welwitschia trees (*Welwitschia mirabilis* Hooker 1863; welwitschia) were highlighted. *Welwitschia mirabilis* is regarded as a living fossil, representing an ancient lineage of gymnosperm plants and it is recognized as a symbol of the Namib Desert biodiversity. This species has a peculiar morphology, being a long-living dwarf tree with only two leaves growing throughout the entire plant life (Roskov et al., 2019). This is also a key species in the Namib ecosystems, where it provides food, water, and refuge for many animal species (Henschel and Seely, 2000). The distribution of *W. mirabilis* is restricted to the central and northern Namib Desert, extending from the Kuiseb River in Namibia to the Nicolau River, north of Namibe, in Angola (Giess, 1969; Kers, 1967). In this area, welwitschia trees occur in four separated sub-ranges, three in western Namibia (Fig. 1A) (Bubenzer et al., 2004) and one in south-western Angola. Bombi (2018) showed that populations living in three Namibian subranges have experienced and will face rather different climatic conditions, and a significant reduction of climatic suitability is expected in the northernmost Namibian subrange under current climate change. In particular, the ongoing rise of temperature can drive the local climate out of the realized niche for the northern populations, thus increasing their extinction risk (Bombi, 2018). Although these findings were potentially important for conservation planning, the study was based on low spatial resolution data available at the national scale, thus limiting its utility at a finer scale for such purposes.

**Figure 1.**
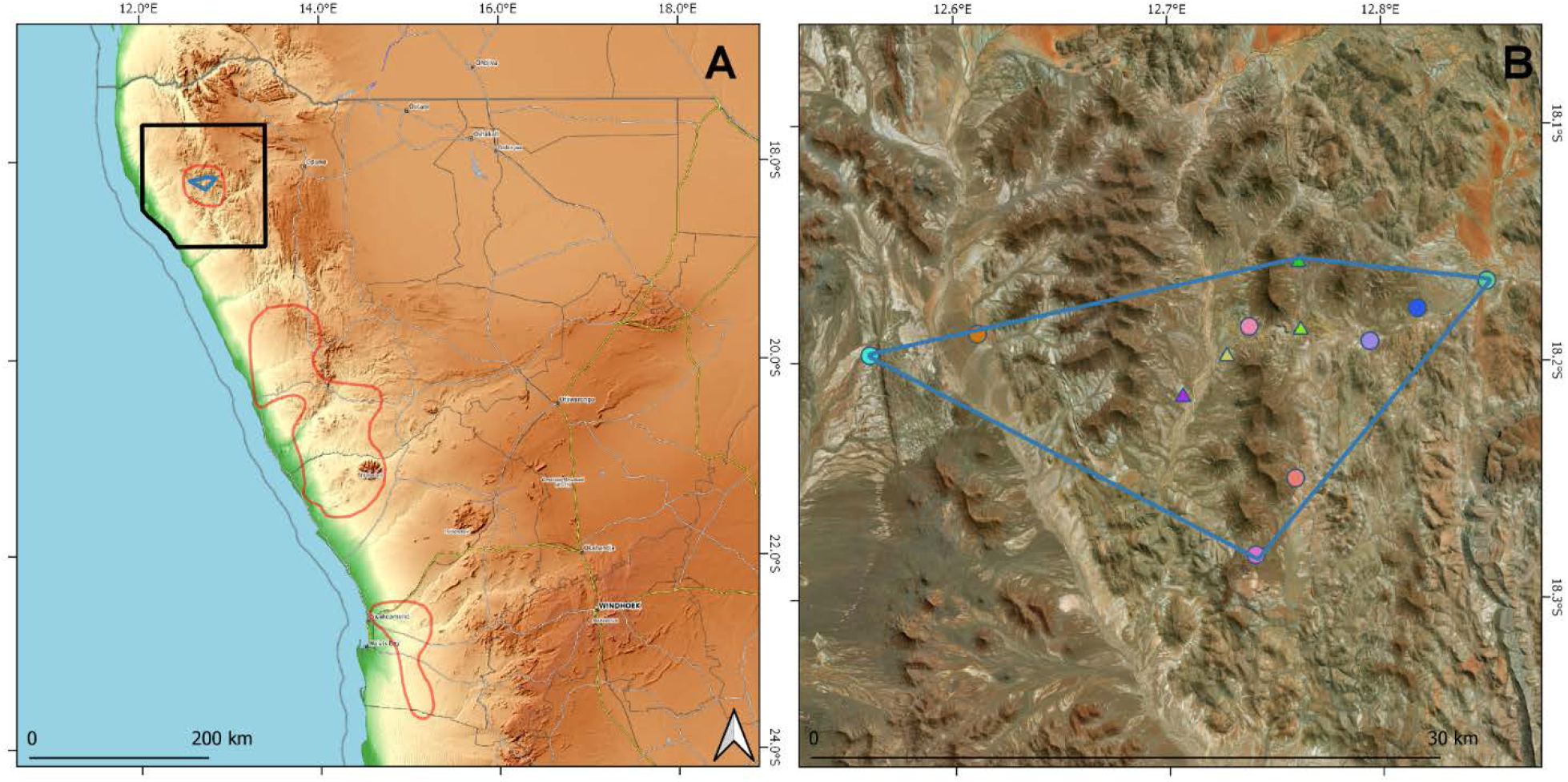
Distribution of *Welwitschia mirabilis* in Namibia and location of the study area (A) and position of detected plant stands inside the observed extent of occurrence (B). In A, the black polygon indicates the study area and the red polygons show the known species distribution. In B, the colored triangles are the plant locations that were known before our study and the colored circles are the new occurrences. In both the maps, the blue polygons represent the boundaries of the observed extent of occurrence in Northern Namibia.

Distribution ranges shifts are well-known responses of species to climate change (Araújo and Rahbek, 2006; Garcia et al., 2014; Parmesan, 2006). These shifts have been generally described as poleward and upward movements of species to track suitable temperature conditions along latitudinal and altitudinal gradients (Hickling et al., 2006; Parmesan et al., 1999; Thomas, 2010). However, in many cases documented geographic patterns of response are complex and do not align with simple latitudinal and altitudinal shifts (Fei et al., 2017). Indeed, the assumption of simple, uni-directional distribution shifts does not account for intricate interactions among temperature, precipitation, and species-specific tolerances and can drive to substantially underestimate the effect of climate change on species distributions (VanDerWal et al., 2012). To overcome these drawbacks, one promising approach is based on the comparison of the species-specific spatial pattern of expected responses, generated by predictive models, with the observed pattern of species response measured in the field with appropriate metrics of population trends (Bombi et al., 2017). This approach can increase our capacity of identifying the footprint of climate change on species dynamics.

The main aim of this study is to verify whether the observed geographic pattern of population conditions of welwitschia trees in northwestern Namibia can be associated to the ongoing climate change. Secondarily, we tested if the same pattern follows a latitudinal or altitudinal gradient in agreement with the assumption of a poleward or upward range shift. More specifically, we want first to validate with field-based data the predictions of potential impacts of climate change on *W. mirabilis*. Second, we want to assess whether the simple assumption of a poleward/upward range shift is suitable for detecting climate change footprints in this case. To do this, we compared the geographic pattern of population conditions, measured in the field, with the expected pattern of response, predicted by ecological niche models. If climate change affects welwitschia populations, we expect worst population conditions in sites where climatic suitability will decrease than in sites where climatic suitability will increase. Moreover, if a poleward/upward range shift is the major response to climate change we can expect a latitudinal or altitudinal trend in the observed patterns of response. Since potential divergent responses to climate change by intraspecific lineages were evidenced (Pearman et al., 2010) and different realized niche were described for each distinct Namibian subranges (Bombi et al., 2017), we focused on populations in the northern subrange and considered them as an independent ecological unit, with its own climatic niche and with its (sub)specific expected response. By verifying our main and secondary hypotheses, we provide documented information to the long-term conservation of *W. mirabilis* and further contribute to the scientific debate on the climate change impacts on biodiversity.

## 2. Materials and methods

### 2.1 Field data collection

During May 2019, we carried out a field expedition in the northernmost Namibian subrange of *W. mirabilis*, as defined by the ‘Digital Atlas of Namibia’ (Bubenzer et al., 2004), in order to obtain information relevant for the species conservation. During the expedition, we spent 10 full days searching for welwitschia trees across the subrange by (1) driving at low speed along the available tracks (more than 330 km) while recording the presence of plants in a ~30 m wide transect on each side of the vehicle, and (2) walking across potentially suitable habitats (more than 65 km). The starting points and spatial extent of our walking transect-based searches were informed by the knowledge of our local team members, who have an intimate knowledge of the area. We are confident that the combination of local knowledge-informed searches and systematic transects extending beyond the known range have allowed us to establish the extent and characteristics of the majority of this sub-range.

During our transects, we collected detailed data on plant location, health condition, reproductive status, and body size. Specifically, we recorded the precise coordinates (using a handheld GPS), the gender, and the presence/absence of cones for almost all the individual plants we observed (just few, unreachable plants were excluded). In sites with a sufficient number of plants, we also measured the stem diameters (minimum and maximum along the two main axes of the stem) and the leaf length, and we recorded the health condition (ranked as dead, poor, average, or good) for a random subset of ~60 plants. We ranked health condition on a four-point scale (dead, poor, average, good) based on leaf color and the general aspect of the plant. Although this is a relatively coarse scale, the brightness of the green color and the ratio of red/brown to green together are a remarkably consistent and accurate indicator of good health condition as measured by photosynthesis efficiency (Shuuya, 2016). The green color of the leaf is associated to the chlorophyll content and the photosynthesis efficiency of the tissues (Menzies et al., 2016; Terashima et al., 2009), which is influenced by environmental stress (Chaves, 1991; Munns et al., 2006). An estimate of health condition such as the above is both a direct reflection of the environmental (including climatic) stress that the plant experiences and an index of the likelihood that its resistance to parasites might be compromised (Mattson and Haack, 1987; Schoeneweiss, 1978). We expected that changes in local climate will be visible in its leaf colour as a quick proxy of plant health.

### 2.2 Observed pattern of response

For each welwitschia stand (defined *a posteriori*, through a GIS-based analysis, as a group of plants separated from the other groups by a distance larger than the intra-group mean distance), we calculated three categories of indicators of population response (derived from plant health, reproductive status, and size) from the field-measured data. For each stand, we calculated the proportion of plants that were dead or in poor, average and good condition. We also calculated the reproductive status (the proportion of plants in the stand that had cones) and the plant size (average stem length, stem width, and leaf length).

### 2.3 Expected pattern of response

We used a spatially explicit approach based on ecological niche modeling to predict the geographic pattern of plant response expected as a consequence of climate change. To do this, we defined our study area as a bounding box three times larger than the latitudinal and longitudinal extent of the previously known subrange of welwitschia in northern Namibia (Bubenzer et al., 2004). Inside this study area, we fitted models on 1000 pseudo-presence/absence points by using climate data from the WorldClim databank (Hijmans et al., 2005) at the spatial resolution of 30 arcsec (about 1 km). In order to control the model-associated uncertainty, we adopted an ensemble forecasting approach (Araújo and New, 2007) in the *R*-based (R Core Team, 2018) *biomod2* Package (Thuiller et al., 2016). In particular, we used Generalized Linear Models (Mccullagh and Nelder, 1989), Generalized Additive Models (Hastie and Tibshirani, 1986), Generalized Boosting Models (Ridgeway, 1999), Classification Tree Analyses (Breiman et al., 1984), Artificial Neural Network (Ripley, 1996), and Random Forest (Breiman, 2001) methods.

Pseudo-presence/absence points were randomly generated across the study area and classified as presence or absence points based on their position inside or outside the species extent of occurrence, generated as a minimum convex polygon from our detailed distribution data. Multicollinearity among predictors was reduced by discarding those with variance inflation factor higher than five (Belsley, 1991). Each independent model was projected into the study area under current climatic conditions and three-fold cross-validated by calculating the true skill statistic (TSS) (Allouche et al., 2006). Finally, we generated a single consensus model of current suitability by calculating the TSS-weighted sum of the independent models. Future climate suitability was predicted by projecting the models into future climatic conditions across the study area. All the available scenarios of future (2050) climate from CMIP5 were utilized for projecting the models. Suitability variation over time was calculated as the difference between future and current suitability and assigned to the observed plant stands on the basis of their location.

### 2.4 Association between observed and expected patterns

For each variable, we tested the linkage between the observed and the expected patterns of responses by adopting a null-model approach (Gotelli and Graves, 1996; Gotelli and Ulrich, 2012; Harvey et al., 1983). First, we quantified the observed correlation between measured values and expected suitability variation in the same sites (*r*_*obs*_) by calculating the Pearson r. Second, we generated in *R* (R Core Team, 2018) (as all the other analyses) 30,000 random permutations of the measured values and calculated the simulated correlation with the expected suitability variation for each permutation (*r*_*sim*_). Third, we calculated the probability of the null hypothesis that the observed correlation was drawn at random from the distribution of the simulated correlations (Gotelli, 2000). Finally, in order to control the familywise error rate due to multiple comparisons, we corrected our *p* values adopting the approach proposed by Benjamini & Hochberg (1995). These corrected *p* values (*p*_*corr*_) measure the level to which the suitability variation (corresponding to the expected response) due to climate change explains the actual responses observed in the different stands.

In addition, we tested whether the observed responses follows a general and simple geographic pattern. In particular, we tested the hypothesis of a latitudinal (equator-to-pole) or altitudinal (low-to-high elevation) range shift. To do this, we adopted the same approach used for testing the linkage between the observed and the model-based expected patterns of responses. In particular, we contrasted each measured variable with the stand latitudes and altitudes. We quantified the correlation between the measured variable and the latitude/altitude, calculated the probability that the observed correlation comes randomly from the simulated correlations after 30,000 random permutations, and corrected our *p* values with the Benjamini & Hochberg (1995) approach. As a result, we obtained an estimation of the extent to which climate change effects can be explained as a simple geographic gradient.

## 3. Results

Overall, we recorded 1330 plants within the known distribution of *W. mirabilis* in northern Namibia. These plants are clustered in 12 distinct stands, which are scattered across the central part of the known range at elevations between 806 and 991 m above sea level (Fig. 1B). Our local team members, who know the area and the species intimately from years of herding goats and cattle, could not point out any more locations than the ones we recorded during the current study. We are thus confident that the plants that we recorded or observed, and the resulting extent of occurrence, represent the majority of plants in this northern-Namibian sub-range. The surface area of each recorded stand varied from 2000 – 825,000 m^2^ and the number of plants per stand varied between four and ⁓400. The extent of occurrence of welwitschia in the area covers about 215 km^2^ and the inter-stand distance varied from 1.8 to 30 km (Fig. 1B). This is a markedly smaller area than the distribution map previously published for the species in northern Namibia (Bubenzer et al., 2004) but a significant improvement of the existing, but unpublished knowledge of plant location in the area.

The available climatic models revealed that the realized thermal niche of *W. mirabilis* in northern Namibia is expected to become completely unavailable within its current extent of occurrence (Fig. 2A). In particular, annual mean temperature within the stands will rise about 1.5 – 2.5 °C, with strong variations among the different scenarios. In contrast, the total annual precipitation will likely remain relatively stable (Fig. 2B), with small reductions or increments forecasted by different scenarios.

**Figure 2.**
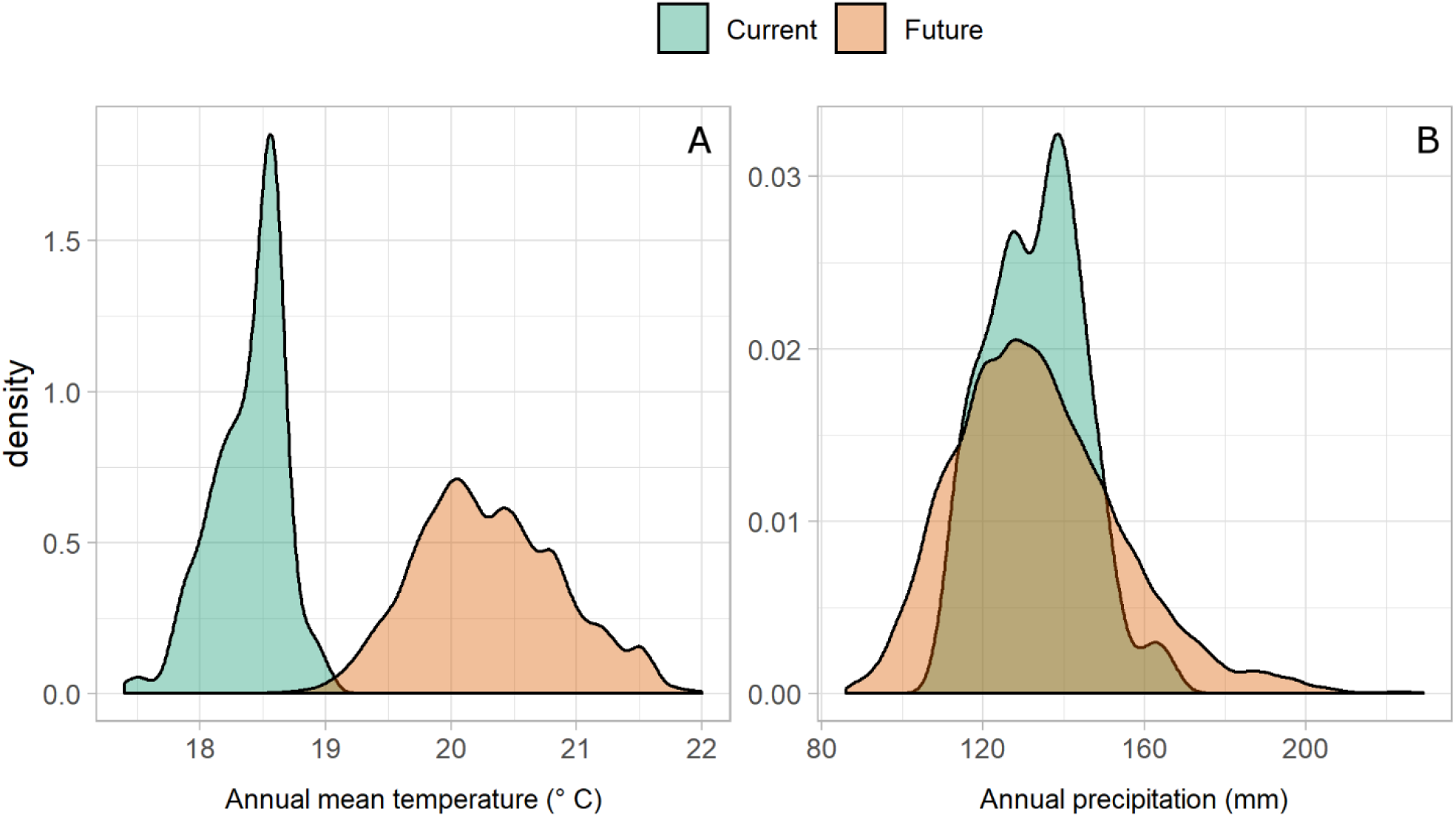
Distribution of current climatic data (i.e. realized climatic niche of *Welwitschia mirabilis*) in the extent of occurrence (in green) and expected future values (in orange) for annual mean temperature (A) and annual precipitation (B).

### 3.1 Observed variation of plant parameters

The most common class of health condition was ‘average’, with 50% of all the plants and a range between 32% and 74% across individual stands being found in this status. Plants in ‘poor’ condition were 32% (range: 11-50%), but only 10% of all plants were in a ‘good’ condition (range: 0-30%). Seven percent of all plants were dead (range: 0-30%) and 56% (range: 10-90%) had cones. Not all individuals could be sexed, but among those that were, 56% were males, with a sex ratio (males/females) ranging between 0.6 – 1.7 across stands. Stem length and stem width were highly variable, ranging from 2 to 100 cm (18.8 ± 14.1 cm; range: 10-33 cm) and from 0.3 and 55 cm (10.3 ± 9.8 cm; range 4.6-22 cm), respectively. Leaf length varied from almost 0 cm (completely browsed plants) up to 93 cm (18.7 ± 13.4; range 11-40 cm).

### 3.2 Expected pattern of species response

The current climatic suitability for *W. mirabilis* is especially high in the eastern half of its extent of occurrence (Fig. 3A). Some areas to the population’s south, as well as a northwest-trending band to the north are also predicted to be highly suitable (Fig. 3A), although plants have never been recorded from these areas before, nor did we find any. Our models further predict that, in the future, the most suitable areas will occur to the northwest of the current extent of occurrence (Fig. 3B). As a result, the plants within in the current extent of occurrence would all experience a reduction of climatic suitability (Fig. 3C) and may thus respond negatively. All the recorded stands are expected to face suitability reductions in the years to come, with variable intensities between almost no reduction and complete reduction. Even those areas that are currently suitable but where the species has not been recorded (Fig. 3A) will similarly experience reductions in suitability (Fig. 3C). In contrast, positive responses may occur in the future suitable area to the northwest of the current extent of occurrence (Fig. 3C).

**Figure 3.**
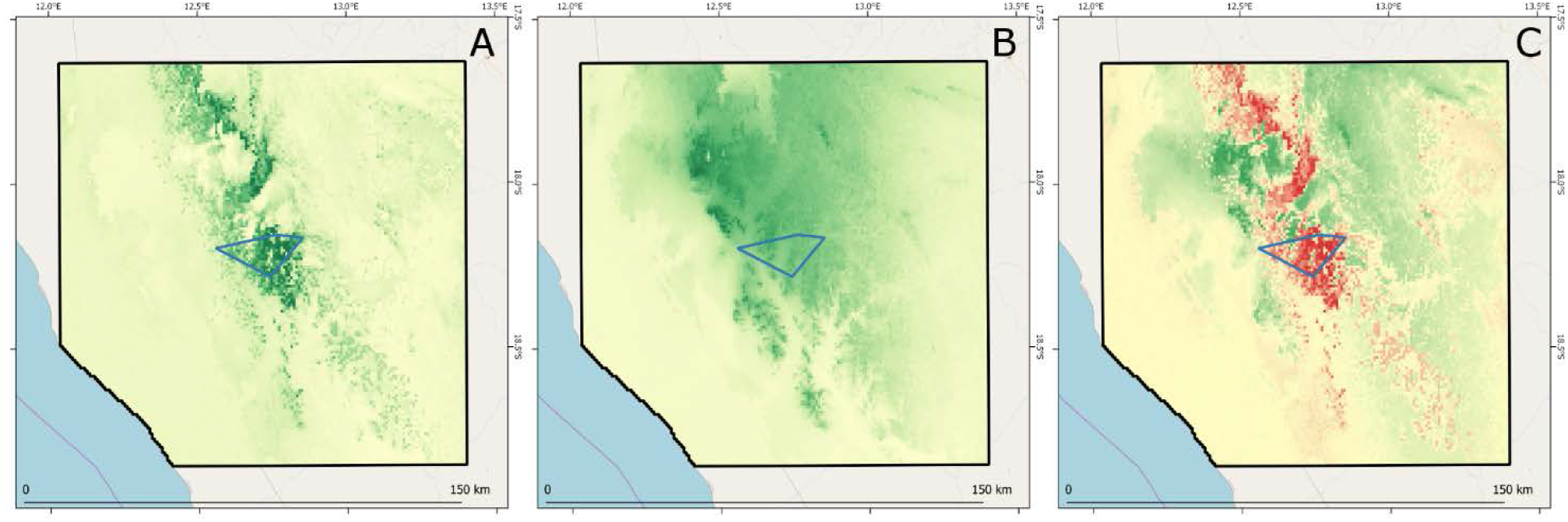
Climatic suitability for *W. mirabilis* in the study area under current (A) and future (B) climatic conditions (green shades indicate growing suitability). Expected suitability variation from climate change (C) (red and green shades indicate negative and positive variations respectively). In all the maps, black polygons indicate the study area and the blue polygons represent the boundaries of the observed extent of occurrence in Northern Namibia.

### 3.3 Observed vs expected patterns

Stronger predicted reductions of climatic suitability in the stand sites are associated with lower plant health condition, fewer plants with cones, and an increased number of dead plants. More specifically, the proportion of plants in poor condition in each stand increases with the reduction of suitability (Fig. 4A). In contrast, the proportion of plants in average and, marginally, of plants in good condition decreases as suitability variation decreases (Fig. 4B and 4C). The proportion of plants with cones (i.e. a proxy of the potential population recruitment) is lower in stands where stronger reductions of climatic suitability are expected (Fig. 4D). At the same time, the proportion of dead plants (i.e. population mortality) is negatively correlated with the expected variation of climatic suitability (Fig. 4E). However, neither the number of plants per stand (i.e. population size) (Fig. 4F) nor plant body size (Fig. 4G, 4H, and 4I) is correlated with the suitability variation.

**Figure 4.**
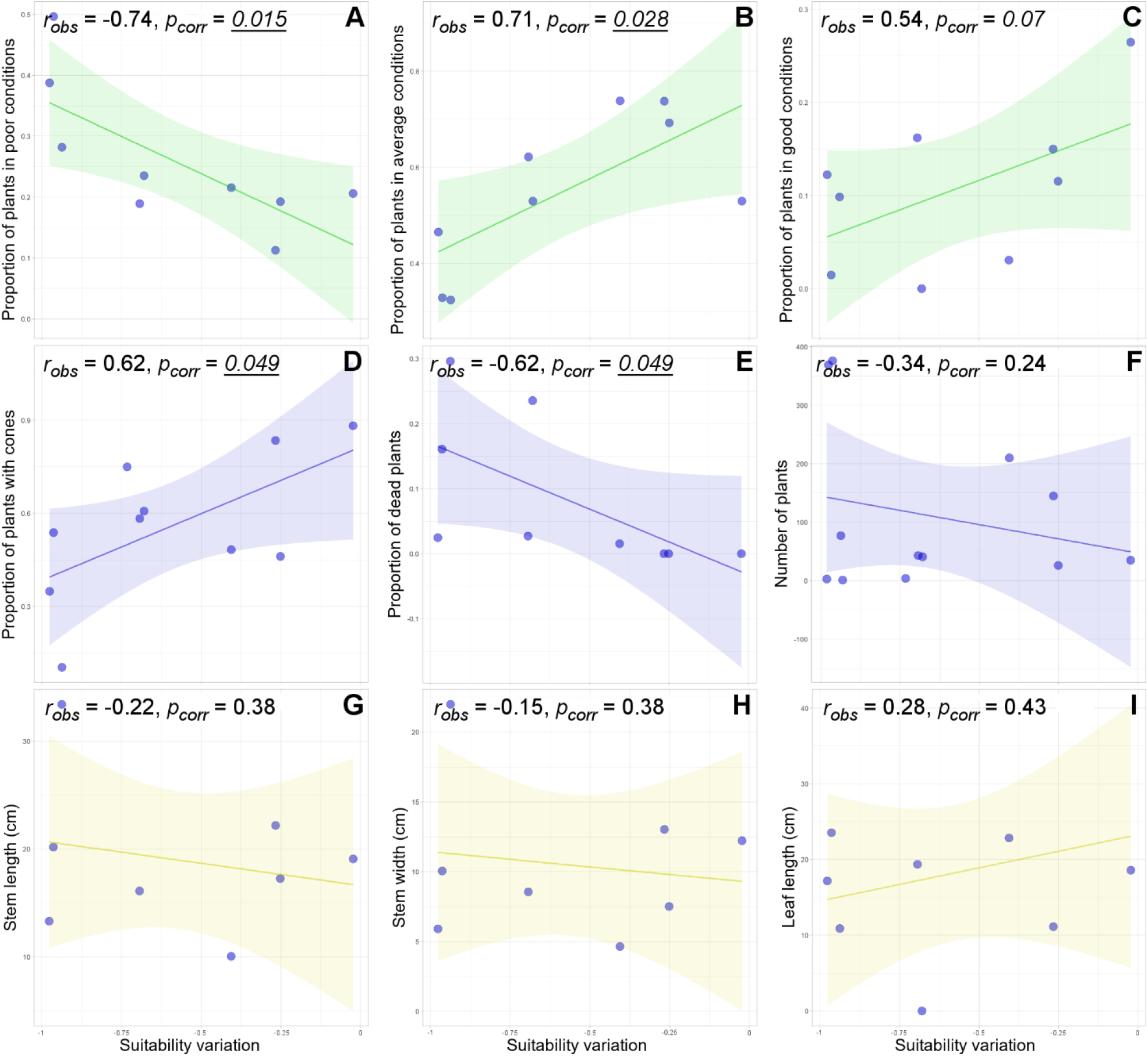
Population features as functions of the expected suitability variation in the stands. In green (first row), features related to plant health condition: proportion of plants in poor (A), average (B), and good (C) conditions. In blue (second row), features related to potential population trends: proportion of plants with cones (D) and of dead plants (E), and number of plants per stand (F). In yellow (third row), features related to plant body size: stem length (G) and width (H), and leaf length (I). In all the plots, blue dots are values for plant stands. Note that suitability variation values (X-axis) are all negative; thus, the reduction of suitability increases from right to left.

The observed geographic pattern of species response does not follow any simple geographic gradient. Indeed, the latitude of welwitschia stands is not correlated with any measured variable (Table 1). Similarly, altitude and the measured variables are not correlated (Table 1). Overall, no latitudinal or altitudinal variation is occurring as a response to climate change.

**Table 1.**
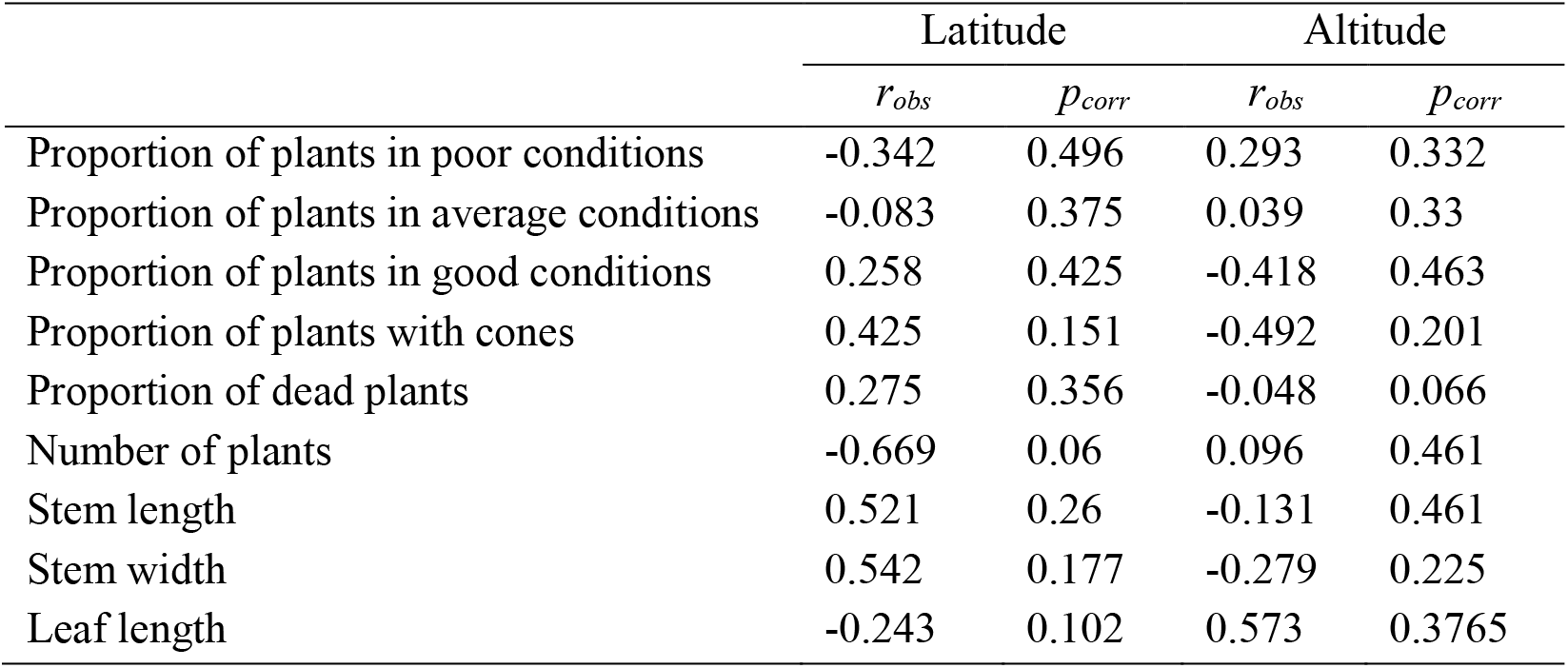
Correlation between the measured variables and the stand latitude and altitude.

|  | Latitude |  | Altitude |  |
| --- | --- | --- | --- | --- |
|  | *r<sub>obs</sub>* | *p<sub>corr</sub>* | *r<sub>obs</sub>* | *p<sub>corr</sub>* |
| Proportion of plants in poor conditions | -0.342 | 0.496 | 0.293 | 0.332 |
| Proportion of plants in average conditions | -0.083 | 0.375 | 0.039 | 0.33 |
| Proportion of plants in good conditions | 0.258 | 0.425 | -0.418 | 0.463 |
| Proportion of plants with cones | 0.425 | 0.151 | -0.492 | 0.201 |
| Proportion of dead plants | 0.275 | 0.356 | -0.048 | 0.066 |
| Number of plants | -0.669 | 0.06 | 0.096 | 0.461 |
| Stem length | 0.521 | 0.26 | -0.131 | 0.461 |
| Stem width | 0.542 | 0.177 | -0.279 | 0.225 |
| Leaf length | -0.243 | 0.102 | 0.573 | 0.3765 |

## 4. Discussion

The results we obtained are coherent with our main hypothesis that the observed pattern of population conditions of welwitschia trees in northern Namibia can be explained as consequences of climate change. Differently, the secondary hypothesis that the geographic pattern of this response to climate change follows a latitudinal or altitudinal gradient is not verified. These outcomes strongly suggest that the ongoing climate change can cause significant alterations to welwitschia populations in the area, can produce important changes (i.e. shifts, contraction) in the local species distribution, and can represent a not negligible threat for the long term conservation of the species. On the other hand, these outcomes also evidence that the potentially serious impact of climate change on this species would be undetected if searched with an approach base on the simple assumption of poleward/upwards range shift.

Inter-stand variations of different population parameters are correlated to change in climatic suitability and can be interpreted as effects of climate change. In this light, the high correlation between the variation of climatic suitability and plant conditions can support a link among climate change, the distribution of plants and the variation of plant health with observed increment of individuals in poor conditions and the reduction of plants in average or good conditions. The loss of climatic suitability can be also put in relationship to the population trends, by affecting recruitment (as suggested by the observed reduction of plants with cones) and mortality (as suggested by the observed increase of dead plants). Even if we measured static parameters of population condition, the pattern of these static measures is coherent with the dynamism of an ongoing range shift from areas currently suitable to areas that will be suitable in the future. Indeed, negative population dynamics, which can be detected as bad population conditions (i.e. scarce plant health, low reproduction potential, high mortality), are typically associated with the trailing edge of a shifting species distribution. The surprisingly clear relationships between observed and expected patterns could indicate one of the very first cases of documented effects of climate change on austral species (Jack et al., 2016).

The higher proportion of dead plants, which suggests an increased mortality, coupled with the lower proportion of plants with cones, which suggests a reduced population recruitment, in areas that suffer stronger effects of climate change, is particularly worrisome because can imply a global negative trend in population size, potentially driving to the local extinction of entire welwitschia stands. In addition, it is worth noting that the linkage between mortality and suitability reduction could be even underestimated by our approach because there are no ways to discriminate plants dead long time ago (i.e. not related to climate change), from recently dead plants. Indeed, the inclusion of old deaths in our dataset could attenuate the climate-related signal. The lack of linkage between suitability variation and plant number could mean that the combination of increased mortality and reduced recruitment has not caused a reduction of population size until now. Similarly, the absence of association between the plant body size and climate change suggests that the altered rates of recruitment and mortality did not modified the age class structure of the stands. Nevertheless, population dynamics of long living organisms can be slow and become evident only in the long term.

As mentioned above, the visual estimation of the plant condition can be considered a rough estimation of chlorophyll content of the leaves and thus of the plant photosynthetic efficiency. Alterations of photosynthesis is a well-known effect of environmental stress (Chaves, 1991; Munns et al., 2006). Heat stress in particular inhibits photosynthesis in tropical and subtropical plants (Larcher, 1995; Salvucci and Crafts-Brandner, 2004). This effect can be stronger in arid environments, where the water shortage can hamper the leaf temperature mitigation (Idso et al., 1982). On the other hand, other studies (Shuuya, 2016) evidenced that, in other populations of *W. mirabilis*, rainfall is followed by an increase in condition. As a result, we can hypothesize that the observed worsening of plant condition is associated to the complex interaction between the significant increment of temperature, which is the main climate alteration expected in the area (Fig. 2), and the constant but limited water availability in the desert environment. Anyway, specifically designed experiments would be needed to tease apart the different possible forces that could cause the observed responses.

Our results confirm the expectation of previous works on the potential impacts of climate change on welwitschia populations in northern Namibia. The study of Bombi (2018), carried out at the national level and at a much coarser spatial resolution, predicted a general reduction of climatic suitability for *W. mirabilis* and suggested potential effects on population recruitment with consequent influences on population structure. At the same time, the author evidenced that living welwitschia trees would have been probably able to cope with the expected climate suitability reduction. On this point, the correlation we observed between increased mortality and predicted influence of climate change would indicate a more worrisome scenario, with a progressive degradation of the plant health and the potential long-term reduction of the population size. This evidence should encourage specific management plans for northern Namibian populations and take into consideration climate change among the conservation issues.

Quantitative data on the plant physiological performances (e.g. leaf growth rate, photosynthesis level, water use efficiency) are required to obtain a more detailed picture of the occurring alterations and to clarify the possible mechanistic linkage with climatic stress. Repeated measures of physiological parameters in different sites would make possible following plant responses over time. The activation of programs for the long-term monitoring of the species in the region would be particularly helpful, allowing critical situations to be detected at early stages and planning effective recovery measures. Obviously, activities of long-term monitoring in this remote area would be difficult and would require the involvement of local communities as well as the provision of significant resources by local and international agencies aimed at the conservation of desert ecosystems in Namibia.

Despite the great interest on *W. mirabilis*, which is considered a living fossil, for its morphological and evolutionary uniqueness (Khoshoo and Ahuja, 1963), and an iconic species of the Namib desert, for its key-role in this ecosystem, several aspects of the species distribution and biology are still to be clarified for a science-based conservation strategy. First, the real level of geographic and genetic isolation of the different subranges should be verified in order to identify intra-specific evolutionary and conservation units. Second, an effort to census and make available the current knowledge on species distribution, demography, and conservation should be undertaken. Indeed, a significant amount of this information is probably pulverized into a plethora of unpublished datasets and field observations. Third, an analysis of the climate change impacts should be extended to the other subranges and a science-based assessment of the conservation status should be made at local and global level. This set of measures could significantly contribute at planning conservation measures for the species that are effective on the long term.

The geographic pattern of response we observed in welwitschia is more complex than the simple poleward/upward shift that was often observed for other species (Parmesan et al., 1999; Root et al., 2003; Thomas, 2010). In the case of *W. mirabilis* populations of northern Namibia, the observed pattern of population conditions, which can represent a response to climate change, follows local contingencies rather than a simple latitudinal or altitudinal trend (Table 1). This could be associated with the small scale of the study but is also in agreement with previous large-scale studies. These studies pointed out that specific responses to climate change can be divergent (Fei et al., 2017) and that assuming a simplified poleward/upward species movement can bring to underestimate climate change impacts (VanDerWal et al., 2012). In our specific case, the linkage between climate change and population conditions, which is suggested by our results, would have been completely undetected with a simplified, but frequently used approach based on the assumption of poleward/upward shifts.

The comparison of the expected pattern of response to climate change, as predicted by suitability modeling, with the observed patterns of population conditions, as measured in the field, appeared as a powerful approach for detecting impacts on wild species. This approach, proposed by Bombi et al. (2017), allowed to indicate climate change as one of the most probable drivers of the geographical variation of population features we observed in the field. This study underlines the importance of considering species responses to climate change as the emergent property of the different effects on individual populations. At a higher biodiversity level, ecosystem responses to climate change can be considered as the emergent property of the effects on individual species. Such a hierarchical relationship provides direction for the application of spatial explicit approaches such as the one used in this study, to multiple species and across diverse ecosystems. In this light, it can be advocated the setting of a large scale program for the identification of sentinel species of climate change effects, which allows to detect, estimate, and follow the climate change impacts on biodiversity and to improve the long-term conservation of species at the ecosystem level.

## Acknowledgements

We wish to thank our Himba guides and friends Riatunga Koruhama and Mavekaumba Tjiposa for the time spent together in front of a campfire and for the willingness to share their gobsmacking knowledge of local environment. We are also grateful to the Okondjombo Communal Conservancy and to the Okondjombo community for their assistance and availability. Karen Nott of IRDNC facilitated access to the area and put us in contact with the community conservancy groups. This study, a sub-project within the Skeleton Coast-Iona Project of the Namibia University of Science and Technology, was authorized through Research Permit RCIV00032018, issued by the National Commission on Research, Science and Technology. Our activities were financially supported by a grant from the Mohamed bin Zayed Species Conservation Fund (Project N 182519816) to P. Bombi.

